# A DltE–DltD–DltX interaction network regulates lipoteichoic acid D-alanylation in *Lactiplantibacillus plantarum* and symbiotic drosophila growth promotion

**DOI:** 10.64898/2026.03.19.713020

**Authors:** Renata C. Matos, Nikos Nikolopoulos, Quentin Perrier, Xavier Robert, Virginie Gueguen-Chaignon, Hiroto Hirayama, François Leulier, Christophe Grangeasse, Yann Guerardel, Stéphanie Ravaud

## Abstract

D-alanylation of teichoic acids is a widespread modification of Gram-positive bacterial cell envelopes that modulates resistance to environmental stresses and host interactions. Although the cytosolic steps of this pathway are well characterized, the extracellular reactions responsible for transferring D-alanine onto teichoic acids remain poorly understood. Here we investigate the role of DltD in the commensal bacterium *Lactiplantibacillus plantarum*. We determined the 2.3 Å crystal structure of the extracellular catalytic domain of DltD, which adopts an SGNH-hydrolase fold with a conserved Ser–His–Asp catalytic triad. Docking analyses with lipoteichoic acids (LTA) fragments suggest that the glycerol-phosphate backbone of LTA is accommodated along a surface groove leading to the catalytic serine, with conserved residues contributing to substrate positioning. Biochemical measurements further reveal direct interactions between DltD, the acyl-carrier protein DltX, and the LTA esterase DltE. The conserved C-terminal motif of DltX binds DltD and is required for efficient D-alanylation and for *L. plantarum*–mediated promotion of *Drosophila* juvenile growth. Together, these findings support a DltX-dependent acyl-transfer mechanism and reveal an interaction network that coordinates LTA D-alanylation in a symbiotic bacterium.

## Introduction

The bacterial cell wall is crucial for defining cell shape, providing mechanical resistance, and facilitating signaling. In firmicutes, a major phylum of Gram-positive bacteria, the cell wall is composed of a thick layer of peptidoglycan - a mesh-like and rigid polymer, which surrounds the cell (1) - and of teichoic acids (TAs). TAs are major anionic glycopolymers either covalently linked to peptidoglycan as wall teichoic acids (WTAs) or anchored to the cytoplasmic membrane as lipoteichoic acids (LTAs) (2–4). These polymers, that are generally synthesized by separated pathways, consist of repeating polyol-phosphate units, such as ribitol-phosphate and glycerol-phosphate, that form intricate networks on the bacterial surface (5). Initially considered as passive structural components, TAs are now recognized as dynamic modulators of bacterial physiology and host interactions, influencing cell morphogenesis, cation homeostasis, autolysin activity, biofilm formation, immune recognition, and adhesion to host tissues (3, 6, 7). Their chemical heterogeneity, generated by glycosylation, phosphocholine substitution or D-alanylation, fine-tunes bacterial adaptation to diverse ecological niches and determines whether bacteria establish symbiosis or trigger infection(7, 8).

Among these modifications, D-alanylation plays a critical role. By reducing the negative charge of TAs, D-alanyl esters impact bacterial resistance to antimicrobial peptides, tolerance to phages, immune evasion, and tissue colonization in pathogens (9–11). In commensals such as *Lactiplantibacillus plantarum* (*Lp*), D-alanylated-lipoteichoic acids (D-Ala-LTAs) were shown to work as symbiotic signals that promote intestinal health and juvenile growth in animal models (12, 13).

D-alanylation is catalyzed by the Dlt machinery, composed of at least four core proteins (10, 14, 15). DltA adenylates D-Ala in the cytosol and transfers it onto the phosphopantetheinyl arm of the carrier protein DltC. The membrane-bound *O*-acyltransferase (MBOAT) DltB subsequently mediates translocation D-Ala on the extracellular side(16, 17). Some models proposed that DltC inserts its D-alanylated arm into the DltB tunnel for direct transfer to TAs. Genetic and biochemical studies also implicate the DltD protein in the final transfer step(3, 15, 18) (Fig. 1A). This raised the possibility that DltD forms a covalent ester intermediate with D-Ala before completing transesterification onto TAs. Yet structural constraints—DltD’s extracellular domain being unable to access the DltB tunnel—challenge this model and suggest that an additional intermediate or accessory factor may be required. A small, conserved motif was shown in *S. aureus* and *S. thermophilus* as the crucial pathway intermediate. This motif is either fused to the DltB protein or constitutes the C-terminal segment of a fifth Dlt protein, DltX composed of a single transmembrane domain (15). In *S. aureus* and *S. thermophilus,* DltX would form a complex with DltB and DltD. The recently published cryo-EM structure of *S. thermophilus* DltB has however further complicated this picture (17). The structure indeed revealed that *S. thermophilus* DltB assembles as a homo-tetramer, and not as a monomer as assumed in previous models. Each protomer binds DltC while maintaining tetrameric organization, and a phosphatidylglycerol molecule was identified in the substrate-binding site as a possible LTA carrier. These findings necessitate a revision of the mechanistic framework for how DltB, DltD, and the accessory protein DltX cooperate to mediate extracellular D-alanylation.

**Figure 1.**
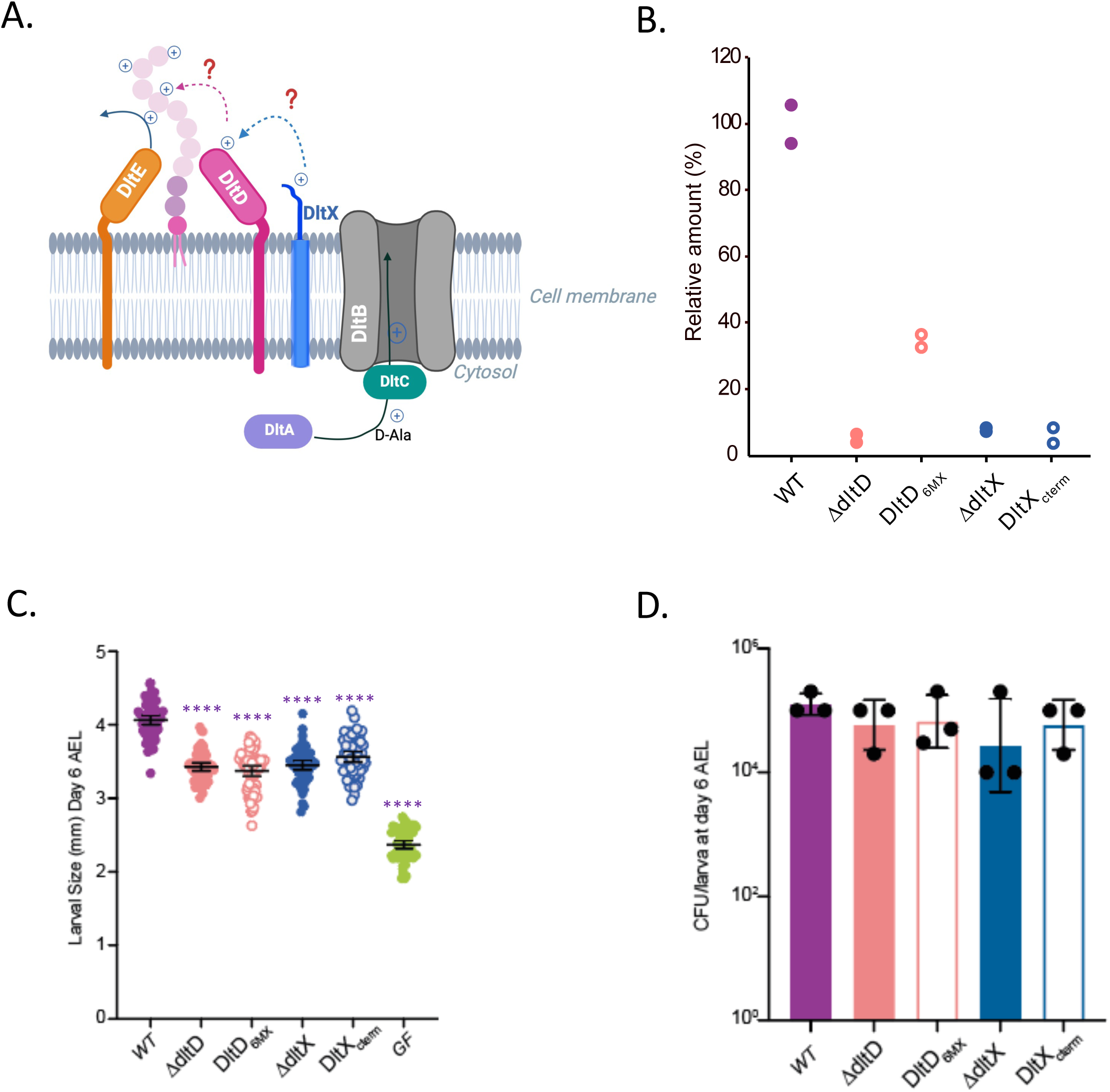
Characterization of *L. plantarum* DltD and DltX mutants in *Drosophila*’s growth. **A,** Schematic representation of the *Lp*Dlt machinery. DltA transfers D-alanine (D-Ala; with a “+” sign) onto the phosphopantetheinyl arm of DltC, which interacts with the MBOAT protein DltB. DltD and DltX are involved in the extracellular steps of the pathway. Their roles are discussed in the paper. DltE removes D-Ala from LTA and may function in coordination with other Dlt components. **B,** Amount of D-Ala released from whole cells of NC8 (WT) and derivative mutants by alkaline hydrolysis and quantified by HPLC after derivatization with Marfey’s reagent. Each value represents a single analysis made on a independent experiment. The data are expressed in % of D-Ala compared to the average value of WT (100%). **C,** Larval longitudinal length after inoculation with strains *Lp^NC8^*, Δ*dltX*, Δ*dltD*, DltX_Δc-term_ and DltD_6M_. Larvae were collected 6 days after association and measured as described in the Methods section. Purple asterisks illustrate statistically significant difference with *Lp^NC8^* larval size; ****: p<0.0001. Center values in the graph represent means and error bars represent 95% CI. Representative graph from one out of three independent experiments. **D,** Bacterial load of *Lp^NC8^* (n=3), Δ*dltX* (n=3), Δ*dltD* (n=3), DltX_Δc-term_ (n=3) and DltD_6M_ (n=3) strains recovered from larvae associated with 10^8^ CFUs after 6 days association. The bars in the graph represent mean and 95% CI. A representative graph from one out of three independent experiments is shown.

In *L. plantarum*, only lipoteichoic acids (LTAs), composed of glycerol-phosphate repeats, are D-alanylated (4, 13). The *dlt* operon contains the core *dltABCD* genes, the additional gene *dltX*, and is further expanded by another gene, *dltE* (Fig. 1A). Importantly, *dltE* is essential for growth promotion in undernourished *Drosophila* by specifically removing D-Ala esters from LTAs, indicating that in *L. plantarum dlt* operon would encode both enzymes that install (DltD) and remove (DltE) D-alanyl modifications (13). This dual activity suggests a regulatory cycle in which the interaction network within the Dlt complex dynamically adjusts LTA structure to optimize host–microbe symbiosis. However, how these components cooperate remains unclear. Whether DltB assembles with DltX and DltD in *L. plantarum*, as reported for *S. aureus* and *S. thermophilus*, remains unresolved. The way DltE integrates into this network is also unknown, as is the precise contribution of DltD, which despite recent advances is still poorly defined. Resolving this question is critical, as it will not only clarify the molecular mechanism underlying TAs modification but also shed light on how this pathway might differ between commensals and pathogens.

Here, we present structural and functional studies of DltD from *L. plantarum*. By integrating biochemical, structural, and genetic approaches, we delineate the role of DltD within the Dlt machinery and uncover how its interplay with other components, DltE and DltX may contribute to symbiotic host–microbe interactions.

## Results

### *Lp*DltD is essential for the TA D-alanylation in *L. plantarum*

*Lp*DltD is a 425-amino-acid protein containing a single transmembrane helix at the N-terminus predicted between residues 7 and 30 and a large domain (residues 31 to 425) predicted to localize extracellularly (Supplementary Fig. 1A). We first aimed to determine whether DltD is individually required for LTA D-alanylation in *Lp*. To address this question, we generated a *ΔdltD* deletion mutant in the *Lp^NC8^* strain using homology-based recombination and quantified the amount of esterified D-alanine (Fig. 1B). Compared with the D-Ala levels measured in the wild-type *Lp^NC8^* strain, the amount of D-Ala released from the *ΔdltD* mutant was nearly undetectable (Fig. 1B). These results confirm that DltD is required for the LTA D-alanylation pathway in *L. plantarum*. Previous work has shown that the *dlt* operon of *L. plantarum* promotes juvenile growth in *Drosophila* under chronic undernutrition (12). To evaluate the specific contribution of *dltD* in this functional context, we assessed the ability of the *ΔdltD* mutant to support the growth of *Drosophila* larvae. Larval size of individuals mono-associated with either the wild-type *Lp^NC8^* strain or the *ΔdltD* mutant was measured six days post-inoculation (Fig. 1C,D). Larvae associated with the *ΔdltD* strain were significantly smaller than those associated with the wild-type strain, despite being colonized equaly by both strains (Fig. 1D). Together, these results demonstrate that DltD is essential for teichoic acid D-alanylation and contributes to the beneficial activity of *L. plantarum* toward its animal host.

### Structure of the extracellular domain of *Lp*DltD

To gain insights into the enzymatic activity of DltD, we cloned and produced the soluble extracellular domain (hereafter referred to as DltD_extra_), which is predicted to contain the catalytic activity (Supplementary Fig. 1B,C). We determined its three-dimensional structure by X-ray crystallography at 2.3 Å resolution (Supplementary Table 1) in the monoclinic space group *C*2. The asymmetric unit contains three molecules that are highly similar (RMSD < 0.2 Å) (Fig. 2A). Analysis of the intermolecular interfaces using PISA yielded a low complex formation significance score (CSS), indicating that these contacts are unlikely to be biologically relevant and most probably arise from crystal packing. The best-defined chain (molecule C) contained 389 out of 402 residues built into the electron density (ranging from 37 to 425). DltD_extra_ adopts a canonical flavodoxin-like fold, consisting of a three-layered α–β–α sandwich with a parallel five-stranded β-sheet (Fig. 2B). Structural comparison with the putative D-alanyl–lipoteichoic acid synthetase from *Streptococcus pneumoniae* (PDB: 3BMA) which served as the molecular replacement model yielded an RMSD of 1.26 Å across 389 aligned residues. Classification using SCOPe and DALI servers revealed that the extracellular domain of DltD belongs to the SGNH-like hydrolase family. Enzymes of this family typically act as esterases and lipases, often targeting the glycerol backbone of lipid substrates. These features support the proposed role of DltD in catalyzing D-Ala esterification on the glycerol phosphate backbone of lipoteichoic acids. In *L*pDltD, the canonical catalytic triad characteristic of SGNH hydrolases is conserved, comprising Ser74, Asp373, and His376. Additional conserved residues Gly104, Gln133, and Trp134 were also identified in the catalytic environment (Fig. 2C and Supplementary Fig. 2).

**Figure 2.**
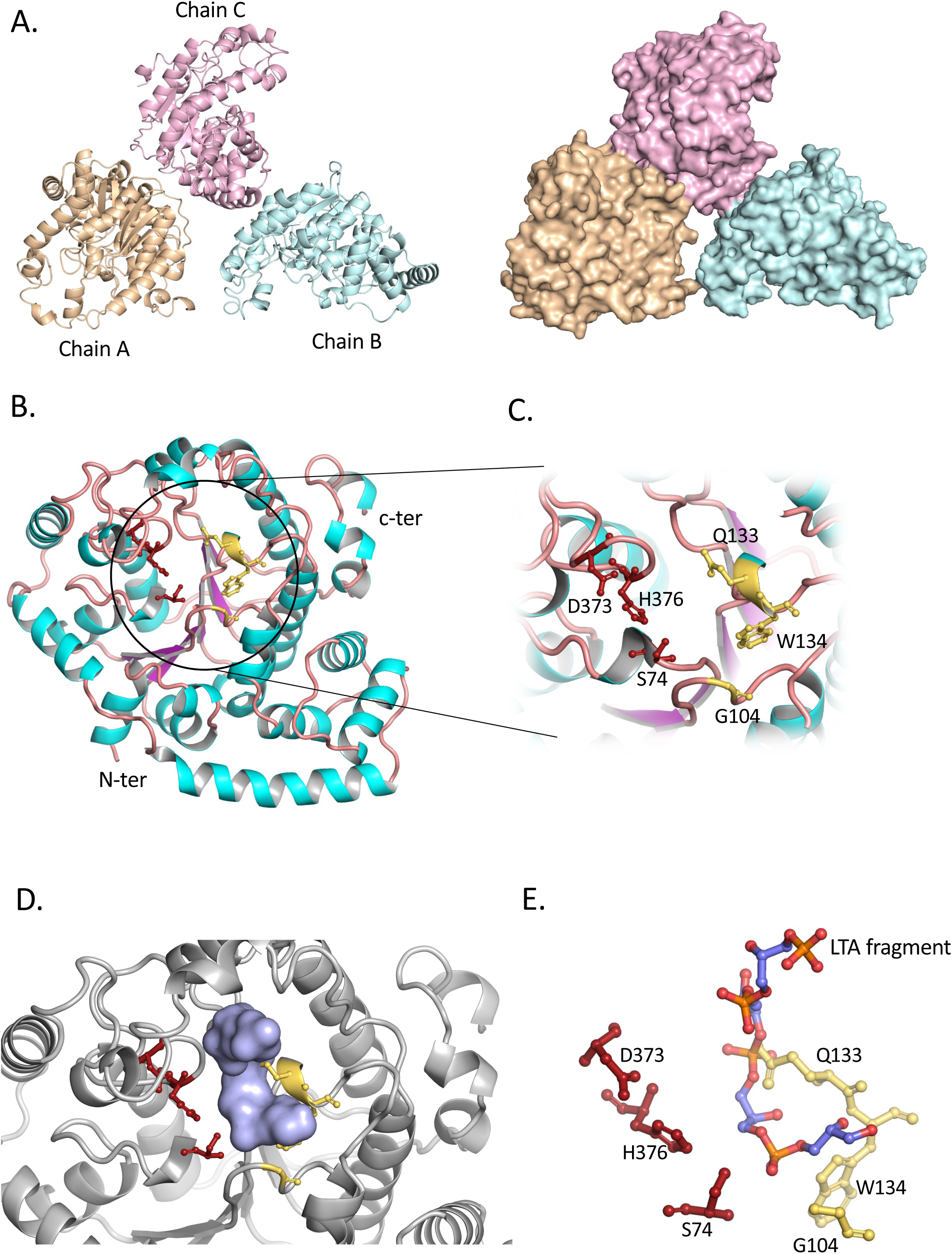
The *L. plantarum* DltD_extra_ structure. **A,** Cartoon and surface representations of the crystal unit cell of the DltD_extra_ structure composed of three chains. **B,** Cartoon representation of the extracellular domain of DltD structure (chain C). The conserved active site residues are shown as red (catalytic triad) and yellow (additional conserved residues) sticks. **C,** Close-up view of the active site. **D,** Close-up on the substrate binding site of DltD with a 4 units LTA fragment docked in the active site. The LTA fragment is skown as a light blue surface. **E,** Close-up of the active-site residues surrounding the 4-unit LTA fragment shown in sticks.

However, the structural basis for substrate recognition by DltD remains unclear. To gain insight into the molecular function of DltD, we performed docking experiments using lipoteichoic acid (LTA) subunits on the experimentally determined apo structure of the extracellular domain of DltD. Docking calculations were carried out with LTA fragments containing three or four glycerol phosphate repeats, corresponding to the LTA composition in *L. plantarum* (Supplementary Fig. 3A,B). Several binding poses with favorable docking scores were obtained for both ligands. In most solutions, the LTA fragments localized to surface pockets of DltD in the vicinity of the catalytic triad (Fig. 2D). Notably, the docked ligands were consistently positioned such that one glycerol phosphate unit was oriented close to the catalytic Ser74 in a geometry compatible with D-alanine transfer (Fig. 2E). The ensemble of docking poses suggests that DltD may accommodate the polymeric substrate in an extended conformation, with successive glycerol phosphate units interacting with positively charged regions of the protein surface. This arrangement could mimic the positioning of the native LTA chain during catalysis and allow sequential processing of individual subunits (Supplementary Fig. 3A,C). Interestingly, the docking results also highlight a potential functional role for several conserved residues—Gly104, Gln133, and Trp134—located near the catalytic site but previously lacking an assigned function. In the predicted complexes, these residues contribute to stabilizing the bound LTA fragments through close contacts, with Gln133 in particular forming interactions with the glycerol phosphate chain (Fig. 2D,E). These observations suggest that conserved residues surrounding the catalytic triad may participate in substrate positioning and stabilization during catalysis.

### DltX is conserved in *Lactiplantibacillus plantarum* and interacts with DltD through its C-terminal motif

The *dlt* operon of *L. plantarum*, and more generally of other *Lactobacillus* species, contains the *dltX* gene, previously implicated in teichoic acid D-alanylation in *B. thuringiensis* and *S. aureus* (15). In *L. plantarum*, we investigated its role by generating a strain lacking *dltX* in the *Lp^NC8^* background. We quantified the level of D-alanylation in the cell envelope and assessed the ability of the mutant to promote *Drosophila* growth. Our results show that *dltX* is required for efficient D-Ala esterification of teichoic acids in the cell envelope (Fig. 1B). Importantly, deletion of *dltX* also abolished the capacity of *L. plantarum* to sustain *Drosophila* juvenile growth, despite colonizing larvae at the same level as the wildtype strain (Fig. 1C,D), indicating that *dltX* is essential not only for teichoic acid D-alanylation but also for the host growth-promoting activity of the bacterium.

We next investigated whether DltX interacts with DltD, as previously shown in *S. aureus* (15). We first used AlphaFold3 (AF3) to model a DltX–DltD complex using the full-length proteins (Fig. 3A and Supplementary Fig. 4A). The predicted structure showed high overall confidence, with consistently high pLDDT scores (Supplementary Fig. 4A). The predicted complex assembly was further supported by strong interface confidence metrics (ipTM = 0.80; pTM = 0.92), indicating a reliable interaction model.

**Figure 3.**
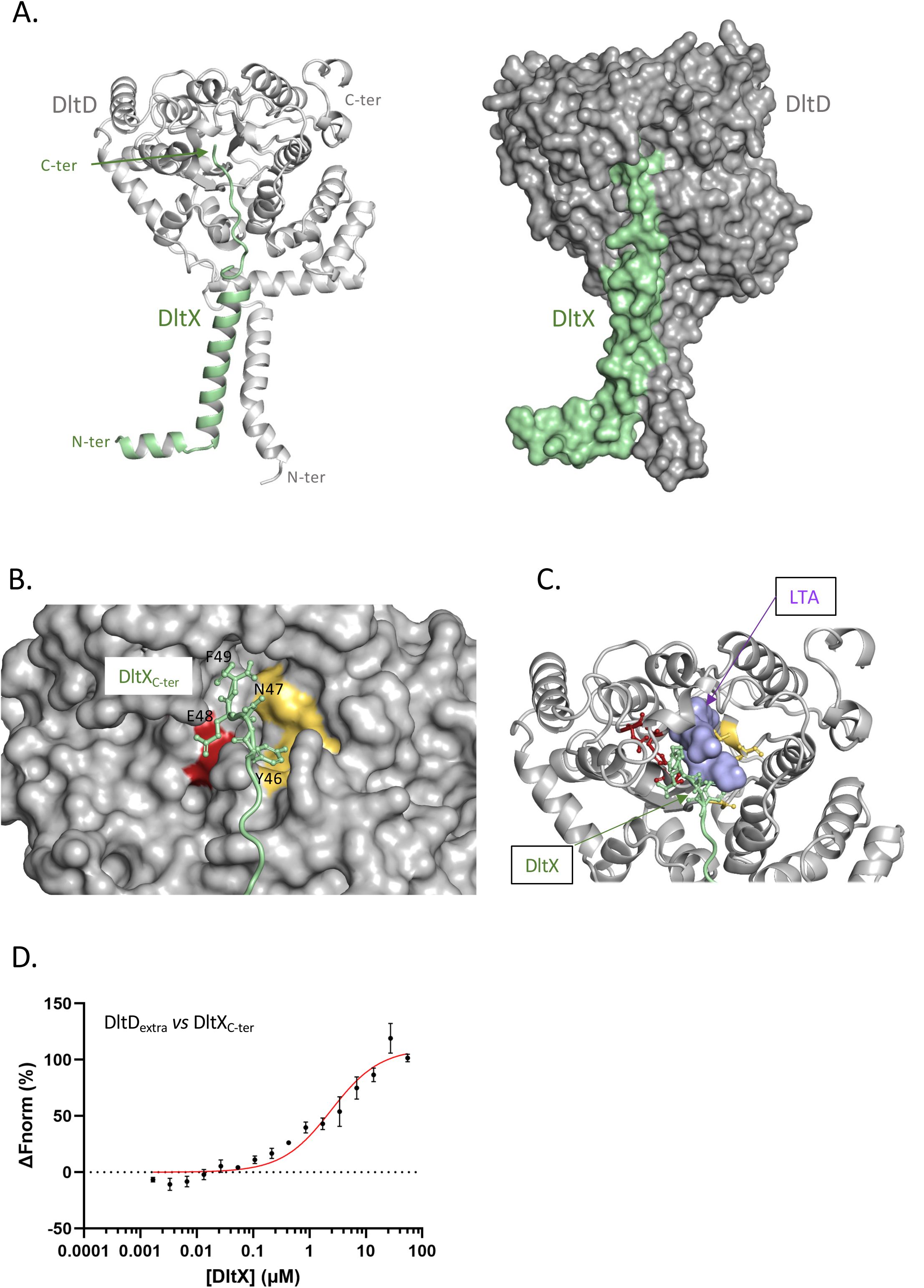
The DltD interaction with DltX. **A,** AlphaFold3 predicted model of the full length DltD-DltX complex as a cartoon and surface representations. DltX is colored in light green and DltD in gray. **B,** Close-up view of the DltD active site, shown as a surface representation, highlighting the predicted position of the C-terminal motif of DltX within the catalytic pocket. The C-terminus of DltX is positioned in proximity to the catalytic residues, which are colored in red and yellow as in Figure 2. **C,** Close-up view of the DltD active site shown as a cartoon representation, with the C-terminal motif of DltX positioned in the catalytic pocket together with a docked LTA fragment containing four glycerol-phosphate units, as in Figure 2. This view highlights that the DltX C-terminus and the LTA fragment bind in the active site. **D,** MST normalized dose–response curves for the binding interaction between the C-terminal peptide of DltX and DltD_extra_ obtained by plotting ΔFnorm against the concentration of the C-terminal peptide of DltX. The binding curves yield a K_d_ of 2.41 ± 0.76 μM. Error bars = s.d.; n = 3.

Analysis of the AF3 model using AlphaBridge (19) revealed two distinct interaction interfaces. One interface involved contacts between the transmembrane α-helices of DltX and DltD, while a second interface was predicted between the C-terminal region of DltX (residues 41-49) and the extracellular domain of DltD (Supplementary Fig. 4B,C). Notably, the last four C-terminal residues of DltX (residues 46–49) were positioned within the catalytic pocket, sandwiched between the catalytic triad of DltD and the predicted binding site of lipoteichoic acid subunits (Fig. 3B,C). This arrangement suggests specific contacts between the C-terminal motif of DltX and the functional surface pocket of DltD_extra_.

To experimentally validate this predicted interaction, we performed microscale thermophoresis (MST) using a synthetic peptide corresponding to the C terminus of *Lp*DltX. These experiments confirmed a direct interaction between the DltX C-terminal peptide and purified DltD_extra_ (Fig. 3D) with a K_d_ of 2.41 ± 0.76 μM.

Sequence alignment highlighted that this C-terminal region of DltX, ^45^FIYNEF^49^, corresponding to the functionally critical motif identified in *S. aureus* (15) is strongly conserved, suggesting an important role for this domain in the D-alanylation pathway (Supplementary Fig. 5). To test this in *L. plantarum*, we generated a *dltX* variant lacking the terminal residues. This variant was defective in D-Ala incorporation and displayed a reduced ability to promote host growth (Fig 1B,C). These results demonstrate that the C-terminus of DltX is essential for the activity of the D-alanylation pathway and for the physiological role of *L. plantarum* in host growth promotion.

Guided by the structural AF3 model and analysis with AlphaBridge and FoldScript (20), we also identified the residues 103, 141–147, 218–221, 273, and 374–375 of the DltD extracellular domain predicted in the interacting interface with the C-terminus of DltX (Supplementary Fig. 4). We then generated a DltD variant (DltD_6M_) carrying six alanine substitutions at predicted interface residues (R103, R141, D218, F221, F273, and T374). Functional analysis of this mutant in *L. plantarum* revealed a reduced D-Ala content in the cell envelope and a decreased ability to promote host growth, indicating that the DltX–DltD interaction contributes to efficient teichoic acid D-alanylation and to the growth-promoting functions of the bacterium (Fig. 1B,C,D).

### Interplay with the *Lp*DltE protein

In *L. plantarum*, DltE was shown to remove D-Ala from LTA. Paradoxically, however, total D-Ala levels remain unchanged in a Δ*dltE* strain or in a strain expressing a catalytically inactive DltE variant (13). These observations indicate that DltE activity may not simply lead to net D-Ala removal from LTA. Instead, they suggest that DltE functions in concert with other components of the Dlt machinery. In this context, DltE may participate in a broader regulatory network that modulates the dynamics of D-Ala incorporation and redistribution along LTA chains, thereby contributing to the overall control of LTA D-alanylation.

Because DltD shares a similar extracellular topology with DltE (Supplementary Fig. 1A, D) and also participates in LTA D-alanylation in *L. plantarum*, we investigated whether the two proteins interact. The extracellular domain of DltE was produced and purified as previously described (13) (Supplementary Fig. 1E,F) and fluorescently labelled to assess binding to DltD_extra_ using MST. MST revealed a direct interaction between the two proteins, with a Kd of 4.7 ± 1.8 μM, consistent with intermediate-affinity binding (Fig. 4). We next attempted to model the DltD–DltE interaction using AF3, both with the full-length sequences and with the isolated extracellular domains. However, neither approach yielded reliable models, as reflected by low interface confidence scores (extracellular domains: ipTM = 0.29, pTM = 0.62; full-length proteins: ipTM = 0.14, pTM = 0.54). These results suggest that the interaction may depend on additional factors not captured in the modelling, such as the presence of LTA or other Dlt components, or a specific oligomeric organization.

**Figure 4.**
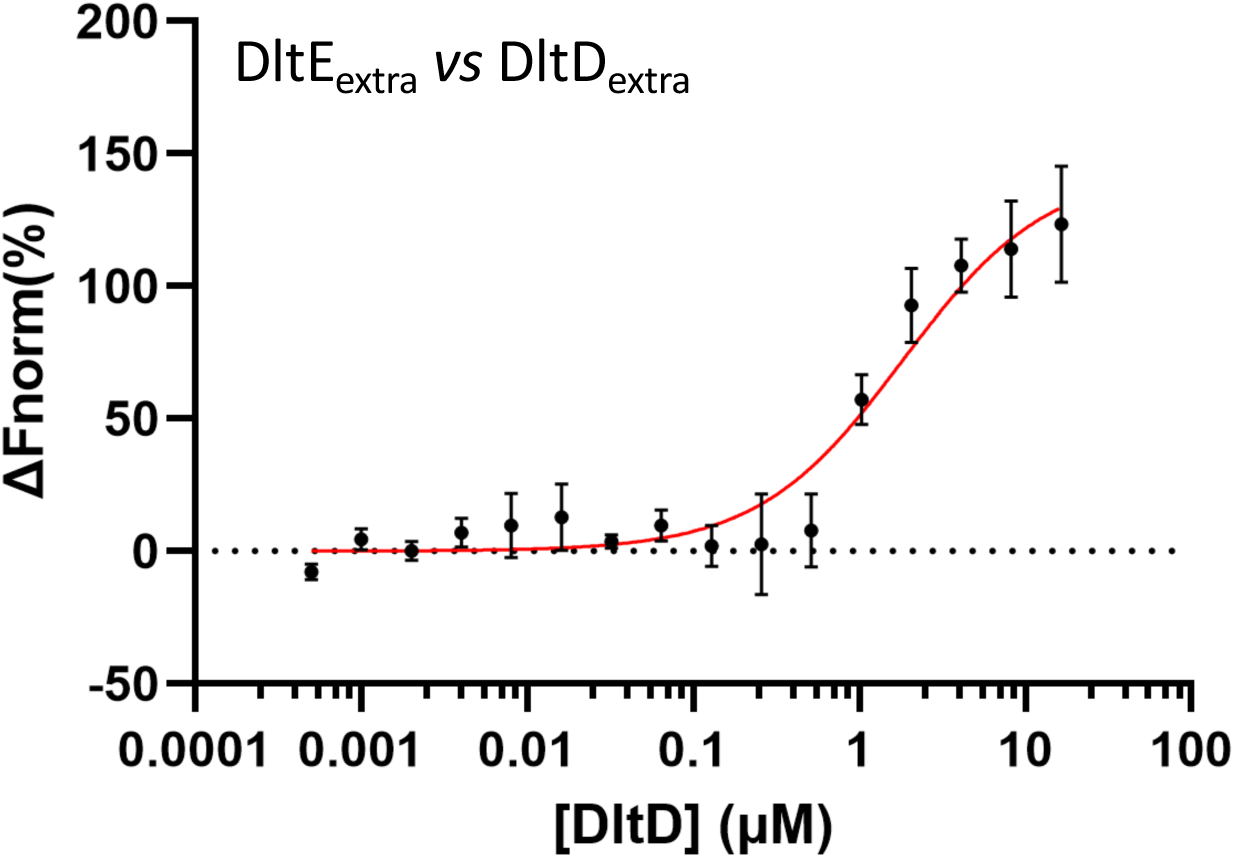
The DltD interaction with DltE. MST normalized dose–response curve for the binding interaction between DltE_extra_ and DltD_extra_ was obtained by plotting ΔFnorm against DltD_extra_ concentration. Plotting of the change in thermophoresis and concomitant fitting of the data yielded a *K_d_* of 1.83 ± 0.55 µM. Error bars = s.d.; n = 3.

Since DltD_extra_ was shown to interact with DltX, we next asked whether DltX could also interact with DltE. We first tested this possibility using a bacterial adenylate cyclase two-hybrid (BACTH) assay. In this system, DltE and DltX were fused to either the T18 or T25 fragments of the catalytic domain of adenylate cyclase (CyaA) from *Bordetella pertussis* (21). A strong interaction between the full-length proteins was detected after 48 h of incubation (Supplementary Fig. 6A). We then examined whether this interaction involved the C-terminal peptide of DltX. MST experiments performed with the extracellular domain of DltE and the C-terminal peptide of DltX confirmed direct binding to DltE (Supplementary Fig. 6B). Consistently, AF3 modelling of a complex between DltE_extra_ and the C-terminal motif of DltX produced a high-confidence model (ipTM = 0.76; pTM = 0.95) (Supplementary Fig. 6C). In this model, the C-terminus of DltX binds within the large cleft of DltE that contains the catalytic residues Ser128 and Lys131 (Fig. 5A,B). Notably, the DltX C-terminal segment occupies a position similar to that of a glycerol-phosphate unit of an LTA fragment observed in the DltE structure (13) (Fig. 5C). This arrangement suggests that the C-terminus of DltX could participate in facilitating D-Ala transfer from the LTA chain bound in the DltE active site.

**Figure 5.**
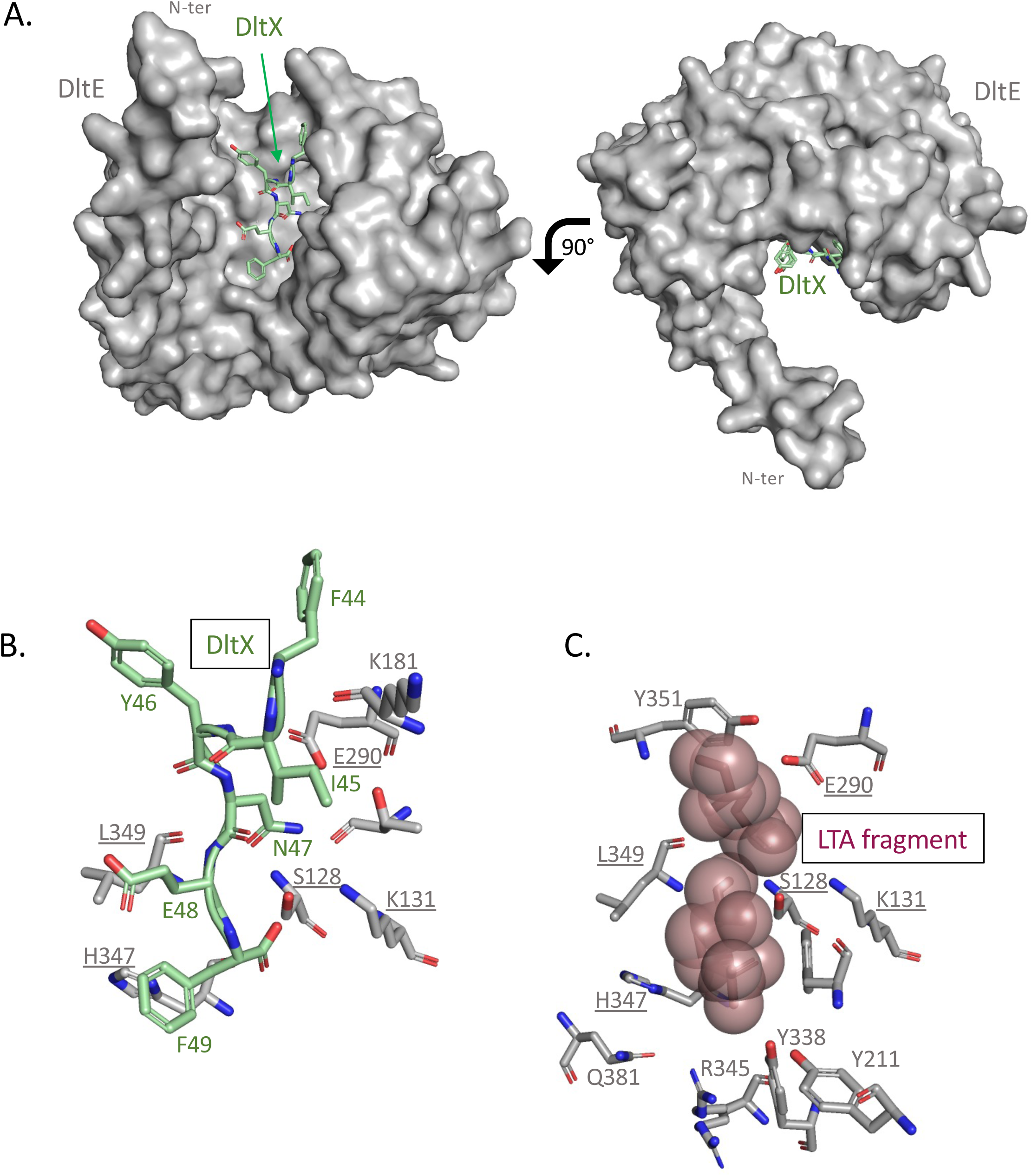
The DltE interaction with DltX. **A,** AF3 predicted model of the complex between DltE_extra_ and the C-terminal motif of DltX consisting of the last 6 residues. DltX, shown in light green sticks binds in the catalytic cleft of DltE shown as a grey surface. **B,** Close-up on the interaction network around the C-terminal motif of DltX modeled in the catalytic cleft of DltE. It involves the catalytic Ser128 and Lys131. This interaction network involves 5 residues that were described in the complex between DltE and a LTA fragment (13) as shown in **C.**

These findings highlight the importance of spatial proximity between DltE and the D-alanylation machinery and point to a key role for the C-terminal region of DltX in mediating these interactions. Taken together, our results suggest that DltX participates in multiple protein–protein interactions within the Dlt system and may function as a molecular connector between DltD and DltE.

## Discussion

In this study, we provide structural, biochemical, and functional evidence that clarifies the role of DltD within the Dlt machinery of *Lp* and situates this enzyme within a conserved DltX-dependent transfer mechanism. Beyond validating a recently proposed acyl-shuttle model, our findings reveal how this conserved enzymatic process is adapted in a commensal bacterium through the integration of an additional regulatory enzyme, DltE, thereby highlighting how a broadly conserved pathway can be tailored to support host–microbe symbiosis.

A central outcome of this work is the confirmation that DltX plays an essential and conserved role in the D-alanylation of LTA. Previous studies in *S. aureus* and *S. thermophilus* established DltX as a small transmembrane protein whose conserved C-terminal motif functions as a transient acyl carrier, receiving D-alanine from the MBOAT protein DltB and transferring it to DltD (15). Our results extend this model to *L. plantarum*, demonstrating that DltX is likewise indispensable for D-alanylation in a commensal context. Importantly, we show that alterations affecting DltX function have direct consequences for host interaction, as disruption of this step impairs the production of D-Ala–modified LTAs that act as symbiotic cues promoting juvenile growth in *Drosophila*. These findings establish DltX not only as an enzymatic intermediate but also as a critical determinant of mutualistic output in a host-associated bacterium.

Our data further demonstrate that the conserved C-terminal region of DltX is functionally central to this process. Perturbation of this short extracytoplasmic motif disrupts the interaction between DltX and DltD, providing direct evidence that the DltX C-terminus is not merely a structural appendage but a key functional interface required for productive D-alanine transfer. This interaction appears essential for the downstream biological effects of D-alanylation, linking precise molecular contacts within the Dlt machinery to the generation of host-active cell envelope signals.

Our results also refine the functional positioning of LTA binding within DltD and suggest that the glycerophosphate backbone of LTA is accommodated along a positively charged surface groove that guides the polymer toward the catalytic serine. In this configuration, conserved residues lining the groove are positioned to stabilize the negatively charged LTA backbone and orient the D-alanine ester linkage for nucleophilic attack. The predicted proximity of these residues to the catalytic triad provides a structural rationale for the strong phenotypes observed upon their substitution and supports a mechanism in which precise substrate positioning, rather than bulk catalysis alone, governs efficient D-alanine transfer. Together, these models explain how DltD can process a large, membrane-anchored polymer while remaining functionally coupled to DltX.

In *L. plantarum*, the DltX–DltD module operates within a more elaborate interaction network that includes DltE, an enzyme absent from canonical *dlt* operons in pathogenic bacteria. We previously established DltE as a D-alanine carboxyesterase that specifically removes D-alanine from LTAs (13), and here we further show that DltE directly interacts with both DltD and DltX. The coexistence of enzymes that install (DltD/X) and remove (DltE) D-alanines within the same operon, together with their direct physical interactions, strongly suggests the existence of a coordinated and potentially regulated cycle controlling the abundance, localization, or timing of D-Ala-LTA presentation at the bacterial surface.

The interplay between DltD, DltX, and DltE raises important questions regarding the spatial and temporal regulation of D-alanylation in commensal bacteria. Our results indicate that productive D-alanine transfer relies on precise coordination between membrane-embedded and extracytoplasmic components and that perturbation of these interactions impacts host-relevant functions. Whether DltE acts constitutively or is itself regulated—by growth phase, nutrient availability, or host-derived signals—remains to be determined. Nevertheless, the direct interactions we observe between DltE and the DltX–DltD module suggest that removal of D-alanine is tightly integrated into the core Dlt machinery rather than occurring as an independent downstream process.

In conclusion, this study advances our understanding of teichoic acid D-alanylation by confirming the conserved role of DltX as an acyl shuttle, clarifying the functional integration of DltD within this mechanism, and revealing the importance of direct interactions between DltD, DltX, and the commensal-specific esterase DltE. Rather than acting as a static biosynthetic endpoint, D-alanylation in *L. plantarum* emerges as a dynamically controlled process shaped by protein–protein interactions within the Dlt machinery. These findings provide a mechanistic basis for understanding how conserved cell envelope modification pathways are adapted in commensal bacteria to support mutualistic host interactions.

## Experimental Procedure

### Bacterial adenylate cyclase two-hybrid assay

The BACTH assay was performed with the BACTH system kit (Euromedex) according to the manufacturer’s protocol (22). Briefly, the Dlt proteins with T18 or T25 were co-transformed and re-streaked on an LB-agar plate supplemented with ampicillin 100 μg/mL, kanamycin 50 μg/mL, isopropyl-β-D-thiogalactopyranoside (IPTG) 0.5 mM, and X-gal 100 μg/mL. Plates were incubated at 30°C in the dark and monitoring of the color shift of the colonies was followed after 12, 24, 48, and 72 h .

### *E. coli* plasmid construction

DNA fragments were amplified by polymerase chain reaction using *L. plantarum* cDNA as a template. The DNA encoding the extracellular domains of wild-type *L. plantarum* DltD and of the mutant DltDS74A were cloned into the NcoI and XhoI sites of the pET-28a(+) vector that expresses proteins fused to a C-terminal hexahistidine (His)_6_ tag. The gene of the DltD-6X harboring R103A, G104A, R141A, D218A, F273A and T374A mutations was synthesized by Twist Bioscience in a pET-28a(+) vector with a C-terminal Hexahistidine (His)_6_ tag. The DNA encoding the extracellular domain of DltE was cloned as described in (13). The peptide NQAKFIYNEF corresponding to the C-terminal sequence of *L. plantarum* DltX was synthetized by the Protein Science Facility platform (PSF, SFR Biosciences).

### Protein production and purification

All the proteins were expressed in *E. coli* BL21 (DE3)-RIPL cells. Cells were grown in Luria-Bertani (LB) medium at 37°C and induced with 0.5mM isopropyl b-D-1-thiogalactopyranoside (IPTG) overnight at 18°C. The cells were then harvested by centrifugation and resuspended in lysis buffer (50mM Tris, pH 7.5, 500mM NaCl, 10% (v/v), Glycerol, 1mM Dithiothreitol (DTT) 0.01 mg/ml Lysozyme, 0.006 mg/ml Dnase/RNase, antiprotease cocktail (CLAPA) composed of Chymostatin 1 µg/mL, Leupeptin 1 µg/mL, Antipain 1 µg/mL, Pepstatin 1 µg/mL, Aprotinin 5 µg/mL). The resuspended cells were disrupted by sonication and centrifuged at 14000 g for 45 min. The extracellular domain of DltD was purified by cation exchange chromatography (HiTrap SP FFcolumn, Cytiva). The starting buffer was composed of 50mM Tris, pH 7.5 and 50mM NaCl and a NaCl gradient (final concentration 1M) was used to elute the protein of interest. The protein eluted at around 500 mM of salt. The eluted fractions were then concentrated and buffer-exchange was performed in a centrifugal filter unit with gel filtration buffer (50mM Tris, pH7.5, 500mM NaCl). The sample was further purified on a Superdex 200 16/600 GL column (GE Healthcare) and eluted with the gel filtration buffer. DltD was finally concentrated but only up to 2.5 mg/mL to avoid precipitation. For DltE, protein production and purification are described in (13).

### Crystallization, data collection and structure determination of DltD_extra_

Crystallization screenings were performed by the sitting-drop vapor-diffusion method using a Mosquito crystallization robot (PSF platform) and the commercial crystallization kits Crystal Screen 1 and 2, PEG/Ion PEG/Ion 2 (Hampton Research). The experiments were performed at 293 K and 70 µL of each precipitant solution were deposited in the reservoir position while 200nL protein solution mixed with 200nL reservoir solution were used for each drop. Crystals appeared after 1 month in the commercial kit solution containing 25% PEG 1K and 0.1 M HEPES pH 7.5. Prior to X-ray analysis, crystals were briefly soaked in a cryoprotectant solution consisting of reservoir solution supplemented with 20% (v/v) ethylene glycol and flash-cooled in liquid nitrogen.

Diffraction data were collected at cryogenic temperature (100 K) on beamline PROXIMA-2A at SOLEIL synchrotron (Gif sur Yvette, France). Data were processed using the XDS package (23). The structures were solved by molecular replacement using Phaser implemented in PHENIX (Adam et al., 2010) using the PDB entry 3BMA as starting model. The structure was refined using iterative rounds of COOT (24) and PHENIX. The quality of the final structure was assessed with MOLPROBITY (25) before deposition at the PDB. The data collection and refinement statistics are presented in Supplementary Table 1.

Sequence alignments and structure images were generated with PyMOL (Schrödinger, LLC), ESPript and ENDscript (26).

### Docking experiments

The 2.3 Å resolution crystal structure of DltD_extra_ and more specifically its best-defined chain C, was used as a target in a protein-ligand docking experiment. LTA fragments containing three or four glycerol phosphate repeats were modelled in 3D using their respective SMILES codes with the Open Babel software (27). The target protein and ligands were then prepared using AutoDockTools v1.5.7: polar hydrogen atoms were added, non-polar hydrogens were merged and Gasteiger partial atomic charges were calculated. Finally, all possible rotatable bonds were assigned to the two LTA fragments. All of the target protein side chains were considered rigid, except for the residues Ser74, Asp373 and His376 of the catalytic triad, as well as the residues Gly104, Gln133 and Trp134 located in the catalytic environment. Two distinct docking experiments were then carried out using the program GNINA v1.2 (28) with the two previously modelled LTA fragments. The search space was a grid box, sized 40×40×34 Å, encompassing the whole DltD_extra_ domain. The following GNINA parameters were employed: num_mc_saved=50, min_rmsd_filter=1.0, cnn=general_default2018_ensemble, cnn_scoring=rescore, cnn_rotation=0, exhaustiveness=40 and num_modes=9. A visual examination of the resulting poses was performed using PyMOL (Schrödinger, LLC).

### Microscale thermophoresis assays

Protein-protein interactions were analyzed by microscale thermophoresis (MST) (29). Buffer of purified and concentrated protein samples was exchanged on a desalting PD-10 column to labeling buffer containing Hepes 25mM pH7.5, NaCl 300mM, Tween20 0.05% (w/v). Proteins were then labeled with NHS red fluorescent dye according to the instructions of the RED-NHS Monolith NT Protein Labeling kit (NanoTemper Technologies GmbH, Munchen, Germany). After a short incubation of target-partner complex, the samples were loaded into MST premium glass capillaries and measurements were performed at 22°C. The assays were repeated three times for each affinity measurement. Data analyses were performed using Nanotemper Analysis software provided by the manufacturer. Data representation was done using Graphpad PRISM 10 software (www.graphpad.com).

### Drosophila diets, stocks and breeding

*Drosophila* stocks were cultured as described in (30). Briefly, flies were kept at 25°C with 12/12 hours dark/light cycles on a yeast/cornmeal medium containing 50 g/L of inactivated yeast. The poor yeast diet was obtained by reducing the amount of inactivated yeast to 7 g/L. Germ-free stocks were established as described in (31). Axenicity was routinely tested by plating serial dilutions of animal lysates on nutrient agar plates. *Drosophila y,w* flies were used as the reference strain in this work.

### Construction of *L. plantarum* strains and growth conditions

Independent markerless deletions on *dltD* and *dltX* genes, and *dltX* c-terminal region of *L. plantarum^NC8^* genome were constructed through homology-based recombination with double-crossing over: as described by (12) (Supplementary Table 2). Briefly, the 5ʹ- and 3ʹ-terminal regions of each region to delete were PCR-amplified with Q5 High-Fidelity 2X Master Mix (NEB) from *L. plantarum^NC8^* chromosomal DNA. Primers contained overlapping regions with pG+host9 (32) to allow for Gibson Assembly. PCR amplifications were made using the following primers: OL01/OL02 and OL03/OL04 (*dltD*), OL07/OL08 and OL09/OL10 (*dltX*), OL21/OL22 and OL23/OL24 (C-terminal region of *dltX*) listed in Supplementary Table 3. The resulting plasmids obtained by Gibson Assembly (NEB) were transformed into *L. plantarum^NC8^* electrocompetent cells and selected at the permissive temperature (28°C) on MRS plates supplemented with 5 µg/mL of erythromycin. Overnight cultures grown under the same conditions were diluted and shifted to the non-permissive temperature (41°C) in the presence of 5 µg/mL of erythromycin to select single crossover integrants. Plasmid excision by a second recombination event was promoted by growing integrants at the permissive temperature without erythromycin. Deletions were confirmed by PCR followed by sequencing.

### knock-in of *dltD* modified version in *L. plantarum*

*L. plantarum^NC8^* strain carrying a modified version of the *dltD* gene was built by knocking-in the modified sequences on *ΔdltD* strain constructed in this study. *dltD* modified sequence (*dltD-6MX*) was synthetized by Twist Bioscience. In order to perform Gibson Assembly (NEB), the overlapping regions were added by PCR with OL15 and OL16 on the plasmid provided by Twist Bioscience. The 5ʹ- and 3ʹ-terminal regions of *dltD* region were PCR-amplified with Q5 High-Fidelity 2X Master Mix (NEB) from *L. plantarum^NC8^* chromosomal DNA using primers OL013/OL14 and OL17/OL18. The 3 fragments were assembled with pG+host9 (32). The resulting plasmids were transformed into *ΔdltD* electrocompetent cells and selected at the permissive temperature (28°C) on MRS plates supplemented with 5 µg/mL of erythromycin. Overnight cultures grown under the same conditions were diluted and shifted to the non-permissive temperature (41°C) in the presence of 5 µg/mL of erythromycin to select single crossover integrants. Plasmid excision by a second recombination event was promoted by growing integrants at the permissive temperature without erythromycin. *dltD* knock-ins were confirmed by PCR followed by sequencing.

### Quantification of D-alanine by HPLC

D-Ala esterified to teichoic acids was detected and quantified by HPLC following its release from intact bacteria. Briefly, D-Ala was released from 10 mg of lyophilized whole heat-inactivated bacteria by mild alkaline hydrolysis with 0.1 N NaOH for 1 hr at 37°C. After neutralization, the extract was incubated with Marfey’s reagent (1-fluoro-2,4-dinitrophenyl-5-L-alanine amide; Sigma) as follows (33). To the sample vial, 100 μL of a 25% (by vol) aqueous solution of triethylamine and 100 μL of a 1 mM solution of Marfey’s reagent in acetone were added and mixed. The reaction was carried out at 40°C for 60 min with gentle shaking in the dark. The reaction mixture was then acidified at room temperature with 20 μL of 2 N HCl and dried under reduced pressure. The residue was dissolved in 50% (by vol) dimethyl sulfoxide before injection. Separation of the amino acid derivatives was performed on a C18 reversed-phase column with a Shimadzu NEXERA UHPLC system with a linear elution gradient of acetonitrile in 20 mM sodium acetate buffer pH 5.0. The eluted compounds were detected by UV absorbance at 340 nm. Relative quantification was achieved by comparison with D-alanine standards.

### Larval size measurements

Axenic adults were put overnight in breeding cages to lay eggs on sterile poor yeast diet. Fresh axenic embryos were collected the next morning and seeded by pools of 40 in tubes containing fly food. 1x10^8^ CFUs or PBS were then inoculated homogenously on the substrate and the eggs. Tubes are incubated at 25°C until larvae collection. Drosophila larvae, 6 days after inoculation, were randomly collected and processed as described by (30). Individual larval longitudinal length was quantified using ImageJ software (34).

### Quantification of bacterial loads

For quantification of bacterial loads, 40 larvae were collected into sterile 100 µm cell strainers and surface-sterilized with 100% ethanol, followed by two rinses in sterile 1x PBS. Larvae were transferred to sterile microcentrifuge tubes and homogenized in 1x PBS with glass beads at 4600 rpm for 30 s (three cycles) using a Precellys 24 tissue homogenizer (Bertin Technologies). Homogenates were serially diluted and plated on MRS agar, and plates were incubated at 28°C overnight to determine colony-forming units (CFUs).

### Statistics

Data representation and analysis of D-alanine quantification and *Drosophila* larval size measurements were performed using Graphpad PRISM 10 software (www.graphpad.com). A total of 3 to 5 replicates were used for all experiments performed in this study in order to ensure representativity and statistical significance. All samples were included in the analysis. Experiments were done without blinding. Two-sided Mann Whitney’s test was applied to perform pairwise statistical analyses between conditions for Drosophila larval size measurements.

## Data availability

Coordinates and structure factors of DltD_extra_ structure have been deposited at wwPDB under the accession code 29GH. Other data supporting the findings of this study are available from the corresponding authors upon reasonable request.

## Supporting information

This article contains supporting information

## Aknowledgments

The authors would like to thank Houssam Akherraz, Nicolas Moreau and Juliette Gonnet for their contributions to the early stages tasks of the project. We are gateful to Céline Freton for help in microscale thermophoresis and to Eric Diesis for the peptide synthesis. We thank the SFR Biosciences (UAR3444/US8) for access to the Protein Science Facility (crystallogenesis robots and peptide synthesis) and the ArthroTools platform (*Drosophila* facility). We acknowledge support on the beamline PROXIMA-2A at SOLEIL synchrotron (Gif sur Yvette, France). This work was funded by the ANR grant ANR-23-CE20-0004-01 awarded to R.C.M and by the collaborative grant ANR-18-CE15-0011 awarded to F.L. and C.G. The laboratories of F.L. and C.G were also supported by the “Fondation pour la Recherche Médicale” (Equipe FRM DEQ20180339196) and by the Fondation Bettencourt-Schueller, respectively.

## Author contributions

Conceptualization: RCM, FL, CG, YG, SR

Methodology: RCM, NN, XR, VGC, ED, YG, SR

Formal analysis: RCM, SR, YG, XR

Investigation: QP, NN, ED, HH, SR, RCM

Writing – original draft: RCM, SR

Writing – review and editing: RCM, NN, QP, XR, VGC, HH, ED, FL, CG, YG, SR

Visualization: RCM, XR, YG, SR

Supervision: RCM, YG, SR, FL, CG

Funding acquisition: RCM, FL, CG

## Supporting Information

### Supporting Figures

**Supplementary Figure 1.**
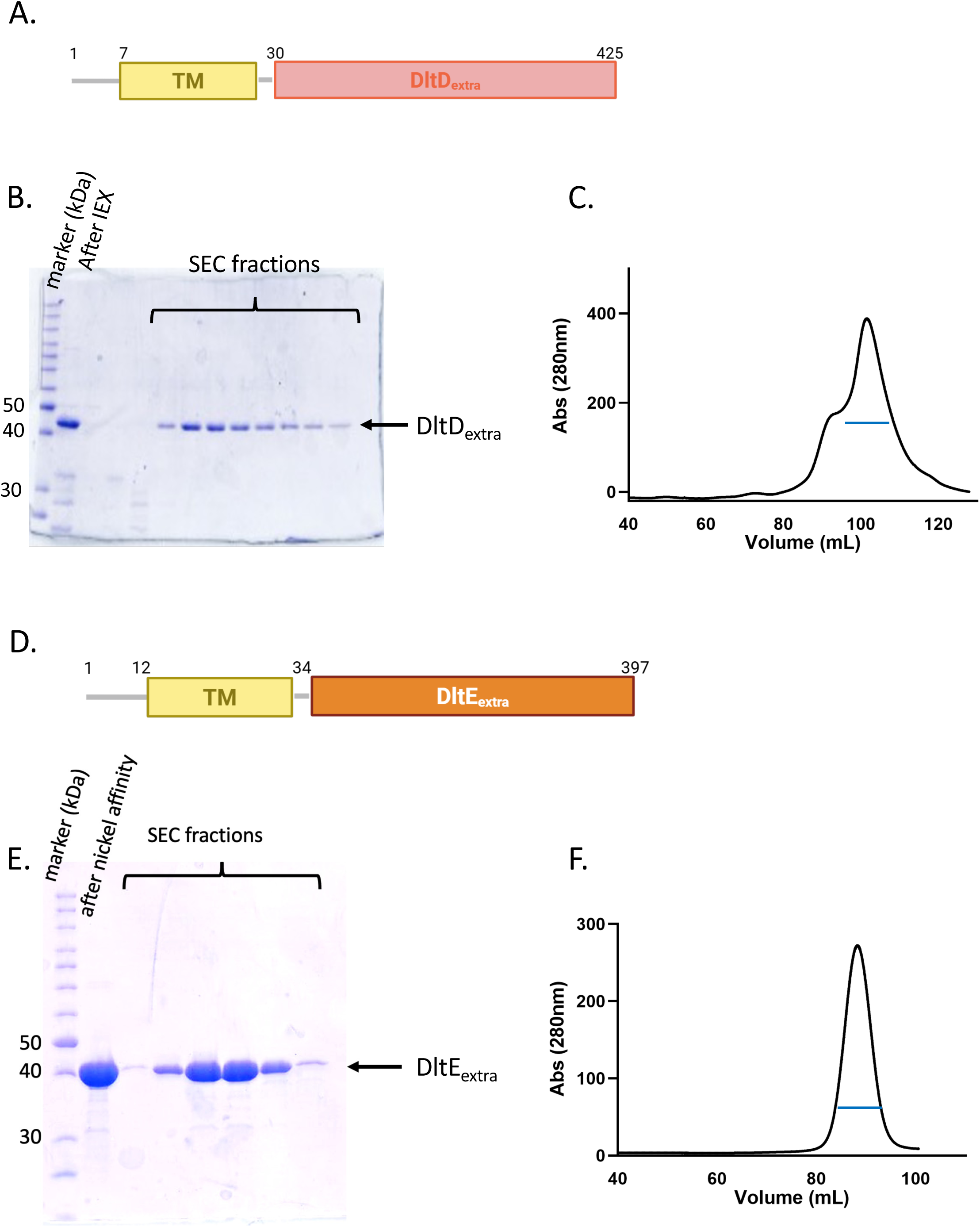
Production of DltD_extra_ and DltE_extra_. **A,** DltD is composed of a transmembrane segment predicted between residues 7 and 30 and a larger C-terminal extracellular region from residues 30 to 425 and named DltD_extra_. **B,** SDS PAGE analysis of the purity of DltD_extra_ after a two-step purification procedure including an ion exchange chromatography (IEX) and a size exclusion chromatography. The band corresponding to DltD_extra_ is shown by an arrow. **C,** Size exclusion chromatography (SEC) and elution profile after injection of the IEX-purified DltD_extra_ on a Superdex 200 10/300 GL. The blue line corresponds to the SEC fractions labeled on the SDS-PAGE. **D,** DltE is composed of a transmembrane segment predicted between residues 12 and 34 and a larger C-terminal extracellular region from residues 34 to 397 and named DltE_extra_. **E**, SDS PAGE analysis of the purity of DltE_extra_ after a two-step purification procedure including a Ni-Affinity and a size exclusion chromatography (13). The band corresponding to DltE_extra_ is shown by an arrow. **F,** Size exclusion chromatography (SEC) and elution profile after injection of the Ni-affinity purified DltE_extra_ on a Superdex 200 10/300 GL. The blue line corresponds to the SEC fractions labeled on the SDS-PAGE.

**Supplementary Figure 2.**
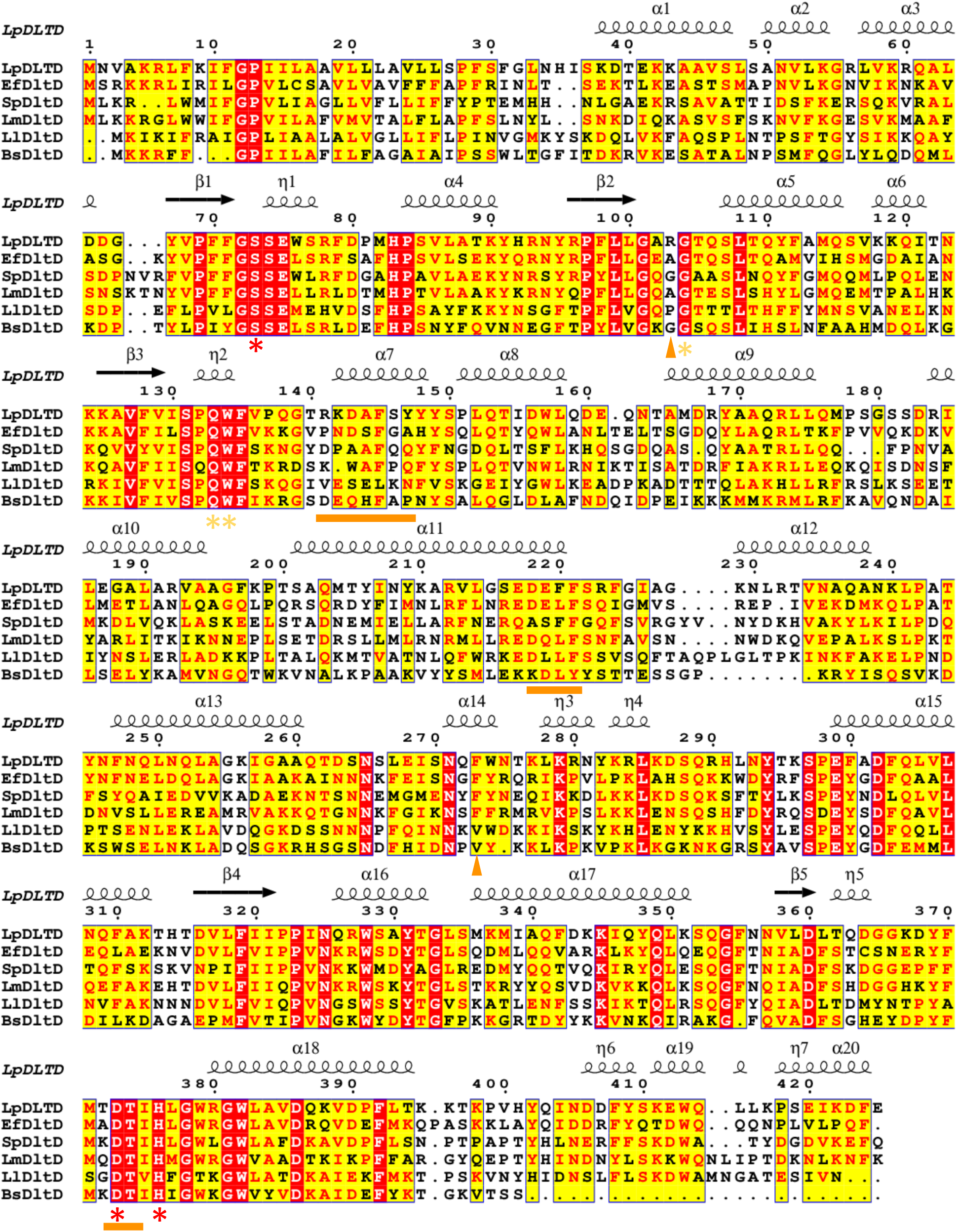
DltD sequence conservation. Sequence alignments between *L. plantarum* DltD and DltD from *Ef, Enterococcus faecalis; Sp, Streptococcus pneumoniae; Ll, Lactococcus lactis; Bs,Bacillus subtilis; Lm,Leuconostoc mesenteroides; Sa, Staphylococcus aureus.* The figure was generated by ESPript (23). The secondary structure extracted from the 3D X-ray structure of DltD_extra_ (this study) is depicted above. The three catalytic residues are marked with a red star while the 3 other conserved residues of the active site are marked with a yellow star. The DltX interacting residues are marked with orange triangles and lines.

**Supplementary Figure 3.**
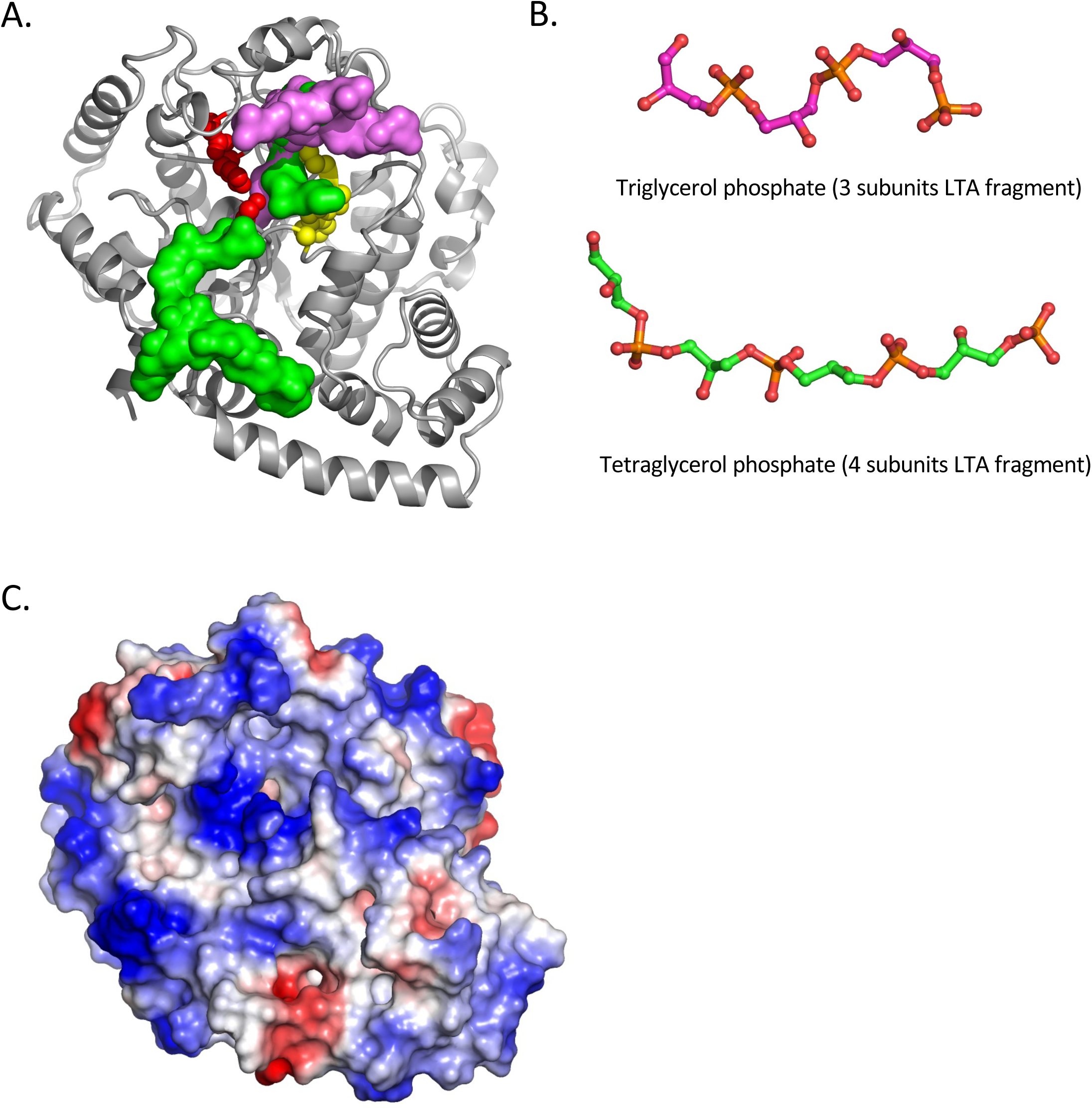
LTA Fragments docked in DltD_extra_ structure. **A,** Docking poses of LTA fragments containing three (in pink) or four (in green) glycerol-phosphate repeats with favorable docking scores mapped onto the crystal structure of DltD_extra_. The protein is shown as a cartoon representation, while the docked ligands are displayed as surfaces. The orientation is the same as in Figure 2, and conserved active-site residues are colored as in Figure 2. The ensemble of docking poses suggests a possible trajectory along the DltD surface that could accommodate the processing of longer LTA chains. **B,** 3D representations of the triglycerol phosphate and tetraglycerol phosphate molecules used in the docking experiments to mimic LTA fragments. **C,** Surface representation of DltD_extra_ colored according to electrostatic potential. The predicted binding poses indicate that LTA fragments preferentially associate with positively charged regions of the protein surface.

**Supplementary Figure 4.**
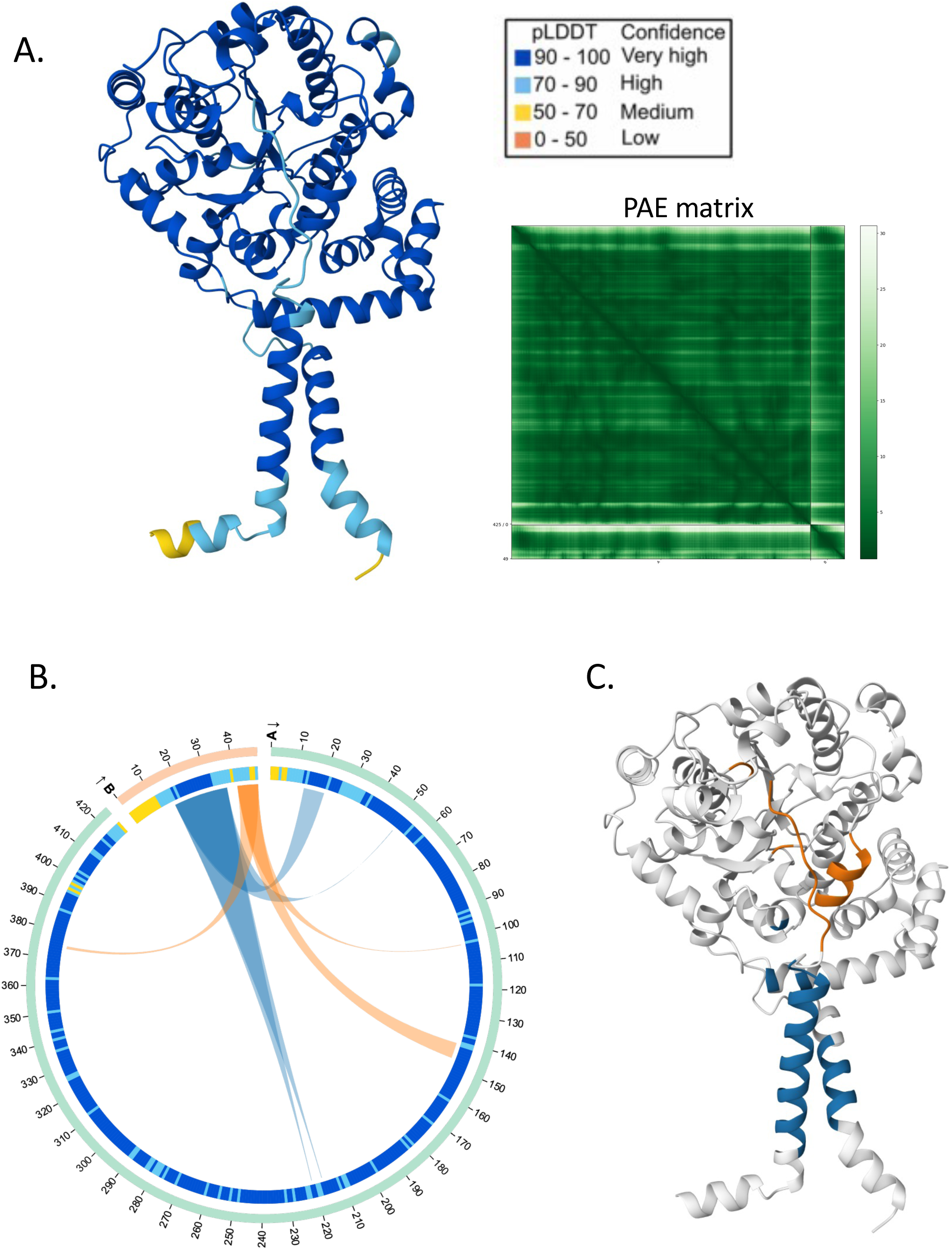
The DltD-DltX structural model. **A,** AF3 model of the DltD-DltX complex and the corresponding PAE matrix. The predicted local Distance Difference Test (plDDT) is a metric of model confidence, with plDDT values > 90 representing very high confidence, 70–90 representing moderate confidence, 50–70 representing low confidence, and < 50 representing very low confidence. **B,** The AlphaBridge plot for this predicted complex. **C,** The two interfaces are mapped on the structure of the complex in blue and orange.

**Supplementary Figure 5.**
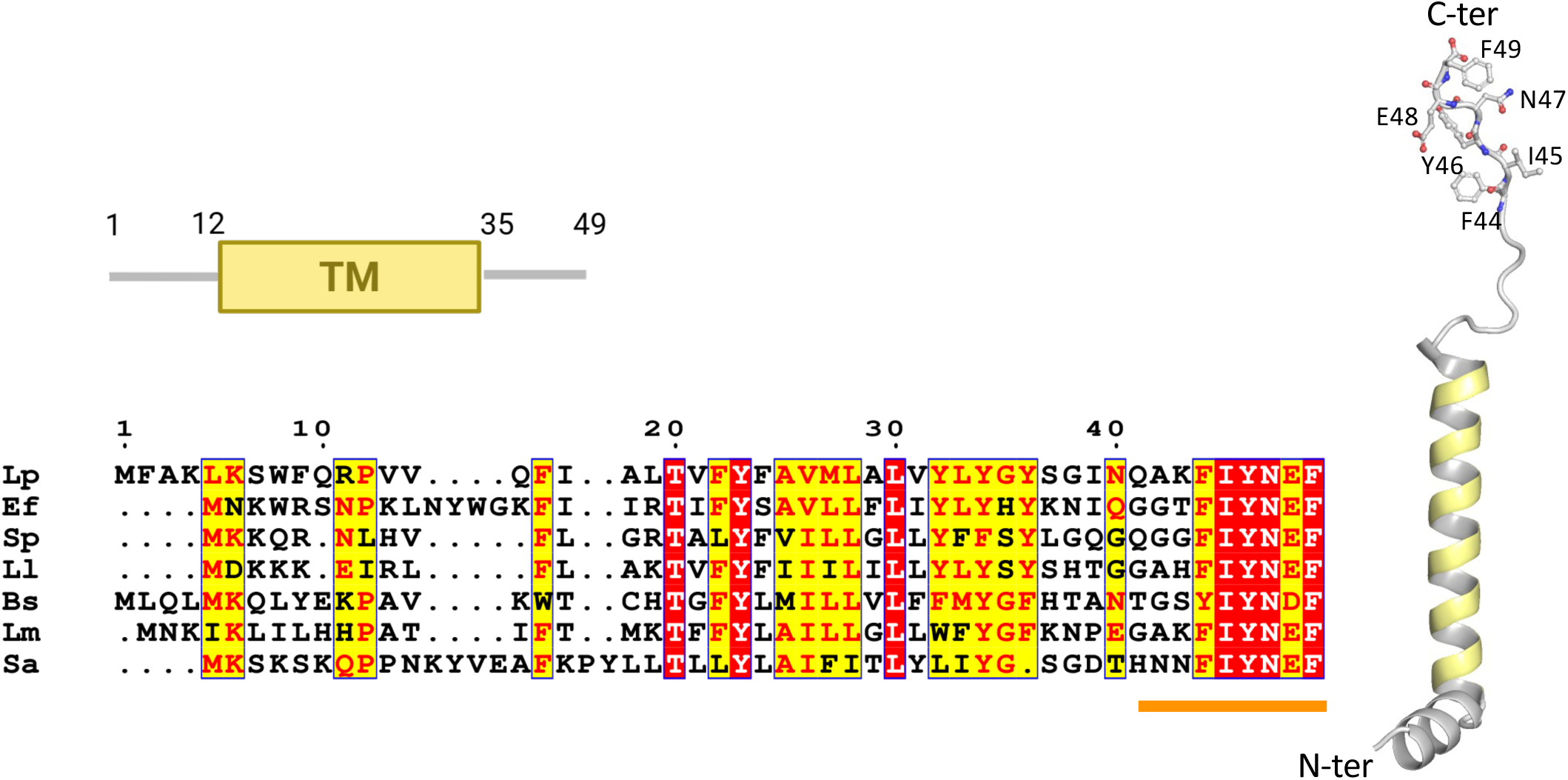
*Lp*DltX sequence and AF3 predicted structure. *Lp*DltX is composed of a transmembrane segment predicted between residues 12 and 35 and a C-terminal extracellular extension of 14 residues. The sequence alignment between *L. plantarum* DltX and DltX from *Ef, Enterococcus faecalis; Sp, Streptococcus pneumoniae; Ll, Lactococcus lactis; Bs,Bacillus subtilis; Lm,Leuconostoc mesenteroides; Sa, Staphylococcus aureus* highlights the sequence conservation of the last 6 residues ^44^FIYNEF^49^. The orange line marks the region predicted to interact with DltD.

**Supplementary Figure 6.**
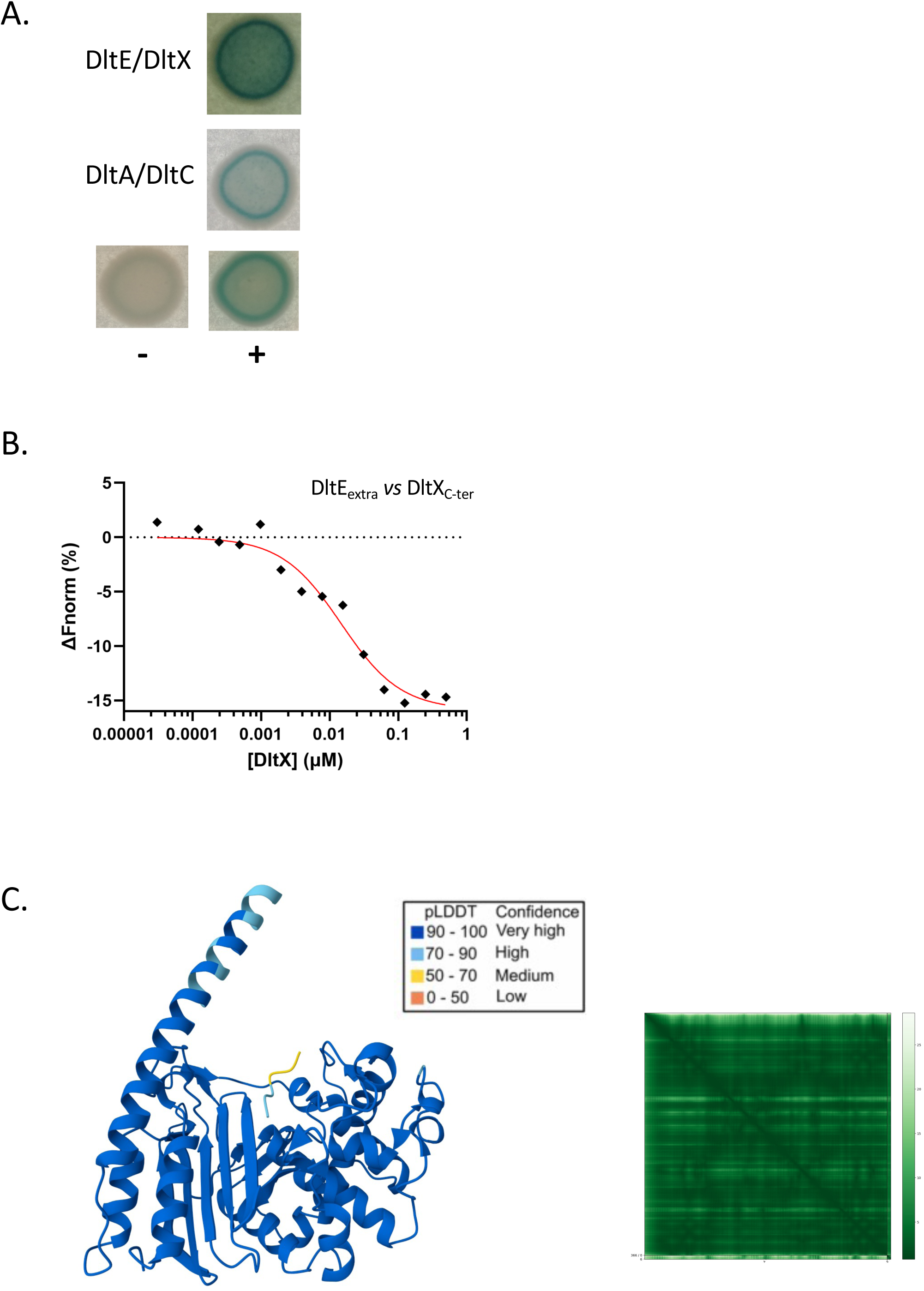
The DltE interaction with DltD and DltX. **A,** The DltE-DltX interaction was assessed by bacterial two-hybrid assay. A blue coloration indicates a positive interaction as depicted by the positive control (+) and the negative control (-) and the comparison with color obtained for the interaction between DltA and DltC that were previously shown to form a complex (14). **B,** MST normalized dose–response curve for the binding interaction between DltE_extra_ and the C-terminus of DltX were obtained by plotting ΔFnorm against the concentration of the C-terminus of DltX. Plotting of the change in thermophoresis and concomitant fitting of the data yielded a *K_d_* in micromolar range. **C,** AF3 model of the DltE_extra_ interaction with the C-terminus of DltX and the corresponding PAE matrix. The predicted local Distance Difference Test (plDDT) is a metric of model confidence, with plDDT values > 90 representing very high confidence, 70–90 representing moderate confidence, 50–70 representing low confidence, and < 50 representing very low confidence.

### Supporting Tables

**Supplementary Table 1.** Data collection and refinement statistics.

|  | **DltD<sub>extra</sub>** |
| --- | --- |
| **Wavelength (Å)** | 0.98013 |
| **Resolution range (Å)** | 44.33 - 2.3 (2.382 – 2.3) |
| **Space group** | C 1 2 1 |
| **Unit cell (Å, °)** | 147.61, 85.33, 110.83,<br>90, 90, 90 |
| **Total reflections** | 438603 (23783) |
| **Unique reflections** | 61272 (6081) |
| **Multiplicity** | 6.9 (5.7) |
| **Completeness (%)** | 99.93 (99.98) |
| **Mean I/sigma(I)** | 10.4 (1.1) |
| **Wilson B-factor** | 42.61 |
| **R-merge** | 0.193 (1.590) |
| **R-meas** | 0.208 (1.746) |
| **R-pim** | 0.078 (0.707) |
| **CC1/2** | 0.995 (0.442) |
| **Reflections used in refinement** | 61272 (2780) |
| **Reflections used for R-free** | 3063 (139) |
| **R-work** | 0.2026 (0.2923) |
| **R-free** | 0.2478 (0.3779) |
| **Number of non-hydrogen atoms** | 9693 |
| Macromolecules | 9421 |
| Ligands | 18 |
| Solvent | 254 |
| **Protein residues** | 1166 |
| **RMS(bonds)** | 0.003 |
| **RMS(angles)** | 0.56 |
| **Ramachandran favored (%)** | 95.34 |
| **Ramachandran allowed (%)** | 3.79 |
| **Ramachandran outliers (%)** | 0.86 |
| **Rotamer outliers (%)** | 0.20 |
| **Clashscore** | 2.46 |
| **Average B-factor** | 48.87 |
| Macromolecules | 48.87 |
| Solvent | 47.86 |
| Ligand | 61.47 |

Statistics for the highest-resolution shell (2.382-2.3 for data collection and 2.4-2.3 for refinement statistics) are reported in parentheses.

**Supplementary Table 2.** Bacterial strains and plasmids used in this study.

| Strain or plasmid | Relevant characteristics | Reference or source |
| --- | --- | --- |
| Strain |  |  |
| *E. coli* |  |  |
| TG1 | supE hsd5h thi (Δlac-proAB) F' (traD36 proAB-lacZΔM15) | Baer et al, 1984 |
| GM1674 | dam-dcm- repA+ | Palmer & Marinus, 1994 |
| *L. plantarum* |  |  |
| NC8 | Isolated from grass silage, plasmid free | Axelsson et al, 2012 |
| ΔdltD | NC8 strain deleted for nc8_1733 | This study |
| DltD-6M | Knock-in of dltD-6M sequence on ΔdltD mutant | This study |
| ΔdltX | NC8 strain deleted for nc8_1737 | This study |
| DltX$_{\Delta$cterm | NC8 strain deleted for dltX c-terminal sequence | This study |
| Plasmids |  |  |
| pG+host9 | Ermr, repATs | Maguin et al, 1996 |

**Supplementary Table 3.**
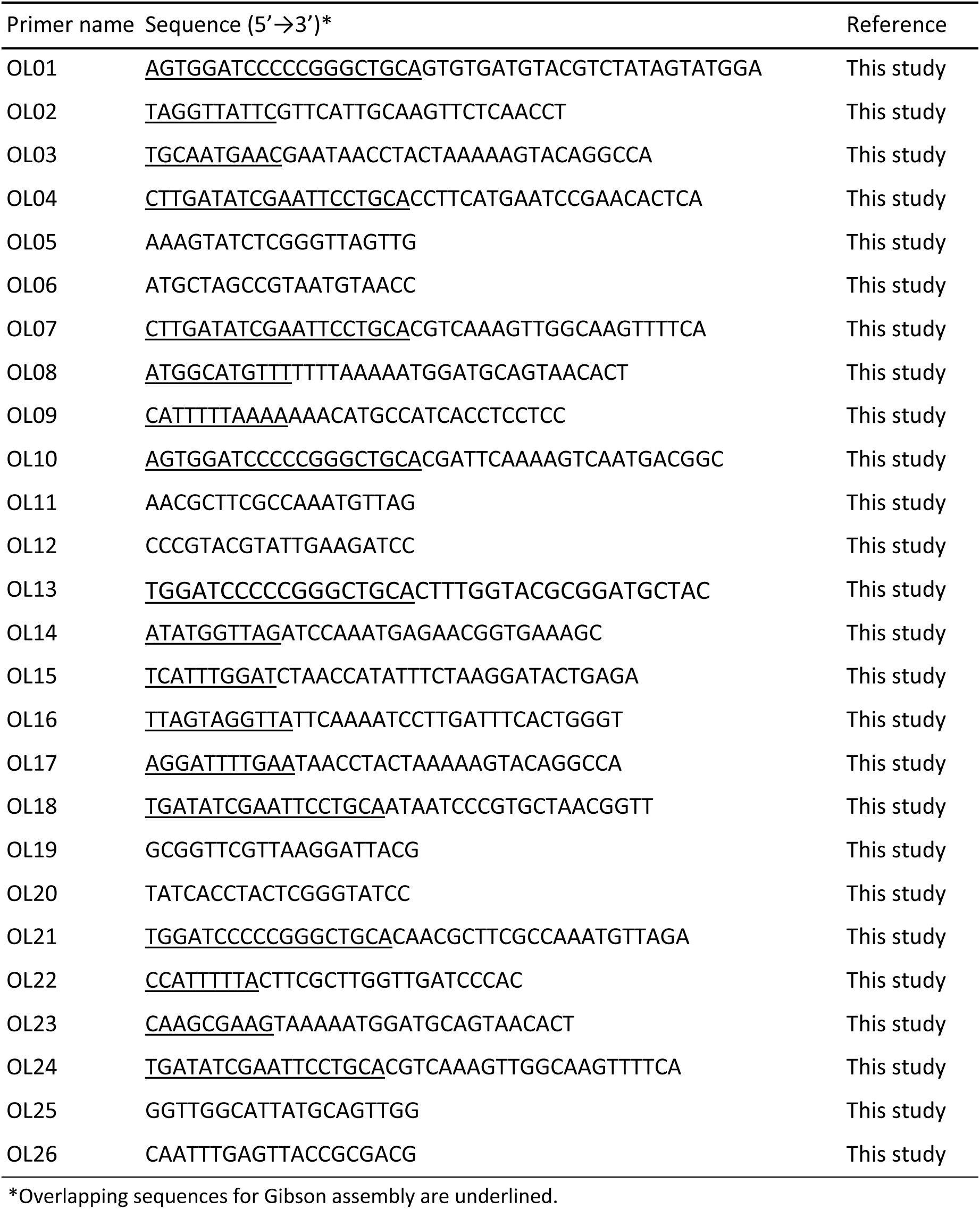
Primers used in this study.

